# Stigma longevity is not a major limiting factor in hybrid wheat seed production

**DOI:** 10.1101/2024.09.13.612789

**Authors:** Marina Millan-Blanquez, James Simmonds, Nicholas Bird, Yann Manes, Cristobal Uauy, Scott A. Boden

## Abstract

Hybrids offer a promising approach to improve crop performance because the progeny are often superior to their parent lines and they outyield inbred varieties. A major challenge in producing hybrid progeny in wheat, however, lies in the low outcrossing rates of the maternal parent. This is often attributed to suboptimal synchronisation of male and female flowering as delayed pollination can result in reproductive failure due to female stigma deterioration. To test this accepted dogma, we examined the seed set capacity of six male sterile (MS) cultivars, each varying in the onset of stigma deterioration. To mimic a hybrid seed production scenario, MS cultivars were grown during two consecutive field seasons, and open pollination was allowed up to 15 days after flowering of the female parent using a blend of seven male fertile cultivars with varying flowering times. Detailed analysis of the temporal and spatial distribution of hybrid seed set along the spike across the six MS cultivars showed that grain production remained remarkably stable during the pollination window tested. These findings suggest sustained receptivity of stigma to pollen across all tested MS cultivars throughout the entire time course. We therefore conclude that stigma longevity does not represent a limiting factor in hybrid wheat seed production, and that breeding efforts should prioritise the study of other female traits, such as enhanced access to airborne pollen.

## INTRODUCTION

In hybrid breeding, two genetically distinct lines are crossbred to produce a first filial generation (F_1_) that typically exhibits superior phenotypic performance compared to its parents, a phenomenon known as heterosis or hybrid vigor. Exploiting heterosis in wheat (*Triticum aestivum*) has been proposed as an alternative for addressing future challenges in food supply (1,2). The use of heterosis promises benefits such as yield increases of 3.5-15% over commercial inbred parents, greater resistance to biotic and abiotic stresses, improved fertiliser-use efficiency, and enhanced yield stability in fluctuating environments (1). However, despite all these advantages and decades of research invested to develop hybrid wheat breeding programmes, hybrids account for less than 1% of the total wheat growing area (3). The limited success of hybrid wheat in the market is due to several factors, including the lack of cost-effective hybrid seed production systems, mainly due to low cross-pollination rates. Another limiting factor is the level of yield heterosis that is perceived by breeders to be too low compared to the productivity of conventional line breeding, with the two-three times higher seed production costs being problematic (4–6).

In self-pollinating or autogamous crops, like wheat, hybrid seed production depends on the unnatural procedure of cross-fertilisation, for which the species are inherently ill-adapted. The discovery of male sterility systems in wheat during the 1960s played a crucial role in facilitating outcrossing, as it provided the means for commercial-scale production of hybrid wheat seed by preventing unwanted self-pollination of the female parent. However, hybrid seed yield on the female parent is still constrained by the structure and development of the wheat flower (5). Consequently, enhancing both female and male parent plants with traits favorable for cross-pollination is essential for efficient hybrid seed production (5). Additionaly, the selection of genetically distinct parents for the purpose of achieving heterosis often results in different flowering times between parental lines. In the case of wheat, the challenge of varying flowering times between the genetically distinct male sterile (MS) line and the pollinator is exacerbated by the short life span of released pollen grains (0.5-3 hours) (7) and the brief duration of stigma receptivity (up to 7 days in moderate temperature and humidity) (8).

Thus, the synchronisation, or ‘nick’, of flowering times between the female and male plants is also essential for the optimisation of seed set in production fields (9). An enhanced understanding of the genes that control pollen availability and viability, and stigma receptivity is crucial, therefore, for improving hybrid seed production.

While the properties and genetic architecture of optimal pollen donor traits are well described (10–12), information about female component traits and their association with improved seed set are comparatively rare (13). Several traits like the stigma length, the openness of the floret or the swelling of the ovary have been described (14–16), but until now, their effect on hybrid seed yield has not been reported.

Stigma development and fertility, which has until recently been a time-consuming and labour intensive trait to measure, has longsince been proposed as a trait to improve hybrid seed set in wheat (17). De Vries summarised studies showing that stigma receptivity lasted between 6 and 13 days depending on the experimental conditions, and described the methods used for calculating receptivity relative to initiation of flowering (18). However, the only data dealing with performance of male sterile plants under open pollination conditions are those reported by Zeven (19).

Zeven determined that the greatest receptivity of male sterile plants relocated for 24 hours to a field of flowering plants happened on the third (1.47 grains per spikelet) and fourth (1.28 grains per spikelet) day after the start of flowering. Thereafter, the receptivity decreased quickly, and on the seventh day practically no more seeds were set. Notably, apart from these studies, there has been limited research dedicated to exploring the influence of female floral traits on hybrid seed set, with most research focused on enhancing pollen-donating qualities (5). This underscores the need for a more comprehensive investigation into the role of stigma receptivity in hybrid seed production.

Previously, we showed that the unpollinated carpel undergoes a well-defined developmental programme that can be used to classify MS cultivars according to the onset and progression of stigma deterioration (17). We hypothesised that MS cultivars with extended stigma longevities could significantly improve the chances of successful pollen germination, fertilisation, and seed set. The aim of this work was to determine the dynamics of stigma development in the female plants of different cultivars and test their influence on hybrid seed production. To achieve this, we selected six MS cultivars showing distinct patterns of stigma development and investigated their contribution to hybrid seed set when open cross-pollination takes place at multiple days after stigma and style development is complete (defined as days after Waddington stage 9.5 (DAW9.5)).

## MATERIALS AND METHODS

### Germplasm

A total of 29 winter male sterile (MS) hexaploid wheat cultivars derived from commercial inbred lines were screened in 2020. Cytoplasmic and nuclear male sterility (CMS and NMS, respectively) systems were used for the generation of the male sterile cultivars. BSS1, BSS2, GSS1 and GSS2 correspond to CMS cultivars provided by Syngenta (Whittlesford, UK) while the remaining are NMS cultivars provided by KWS (Thriplow, UK). Principal component analysis (PCA) and hierarchical clustering analyses were conducted using stigma area data to select six MS cultivars (24485, 24512, 24526, 24522, BSS1, and GSS1) with contrasting patterns of growth and deterioration. These cultivars were further analysed during the 2021 and 2022 field seasons.

For the crossing plots, we used a mixture of pollen donors including seven different male fertile cultivars (Nirvana, Elysee, Piko, Poros, Creator, Quartz and Stava). The selection of these pollen donors was based on visual anther extrusion scores greater than 1.5 (0 = no anthers extruded; 3 = maximum anther extrusion) and distinct flowering times to ensure good pollen availability throughout the time courses (Supplementary Table S1). To avoid unwanted cross-pollination, sterile rye (*Secale cereale*) barriers were grown surrounding the male sterile and crossing plots (e.g., Supplementary Figure S1).

### Experimental design and sampling

Field experiments were conducted during the 2019/2020, 2020/2021 and 2021/2022 growing seasons at John Innes Centre Church Farm (Bawburgh, UK; 52°37’50.7” N 1°10’39.7” E). Plants were grown in a randomised complete block design (RCBD) with two replicates (1 m^2^ plots) per MS cultivar in 2020 (Supplementary Figure S1), and three replicates in 2021 and 2022 for carpel development time course. Hybrid production plots were also grown in RCBD with three replicates in 2021 and 2022 (Supplementary Figure S2, S3). Meteorological data were obtained from data loggers (EasyLog USB, Lascar Electronics; and Tinytag Plus 2) placed next to the experimental plots at 50 cm height. Temperature and relative humidity were measured every hour throughout the experiments (Supplementary Figure S4).

### Carpel development time course

For the carpel development time course experiments, we followed the approach described in (17). Briefly, main spikes were tagged when carpels in florets 1 and 2 of central spikelets reached Waddington stage 9.5 (W9.5). At the time of sampling, we cut individual tillers (two tillers per plot and time point) and transported them in water to the laboratory for carpel dissection. On average, five carpels from central spikelets (floret 1 and 2) were stored in freshly made 95% ethanol and absolute acetic acid (75% v/v) fixative solution and kept at 4°C until image acquisition.

#### Open pollination time course

To control the timing of pollen flow between neighbouring male and MS plants, male and MS plants were separated by isolation walls (1.5 m height protection fleece, LBS Horticulture, Lancashire, UK) in 2021 and pollination bags in 2022 (catalogue No. 2D.4-1W, PBS International, Scarborough, UK), which covered the MS plants before ear emergence (Supplementary Figure S2, S3). At the time of pollination, we removed the walls or bags (depending on the year) allowing cross-pollination to take place from that point onwards. In 2021, the hybrid seed set time course spanned four time points (from 0 to 15 DAW9.5), and six time points (from 0 to 13 DAW9.5) in 2022. Notably, these time points exhibited slight variations among MS cultivars due to divergent phenology, with BSS1 and GSS1 reaching W9.5 later and 24522 reaching W9.5 earlier compared to other MS cultivars.

To determine the level of unwanted cross pollination and level of sterility of MS plants, we included negative control plots in which isolation measures (walls or bags) remained in place throughout the experiment. In 2022, we introduced positive control plots, allowing unrestricted pollen flow (i.e., no bags). It is worth noting that the positive control plots essentially represent the first time point of cross-pollination since, in both cases, pollination was feasible from W9.5 onwards. Each crossing plot consisted of two rows of MS plants flanked by two rows of male fertile plants on either side with 0.17 m spacing between rows (inset in Supplementary Figure S2).

To maximise pollen flow, crossing plots were sown downwind of the main wind direction. At full plant maturity, we hand-harvested between five and ten main spikes per hybrid production plot and 50 spikes from control plots for subsequent hybrid seed set assessments.

### Hybridity test

A set of 93 putative F_1_ seeds were randomly sampled from the 2021 and 2022 field trials for each MS cultivar to undergo hybridity testing. Additionally, 40 seeds from each of the pollen donors were included in the analysis (except for Poros, for which 32 seeds were tested). KWS conducted DNA extraction and genotyping, employing a panel of 23 single nucleotide polymorphisms (SNP) markers. Markers were selected to cover genomic regions showing high levels of diversity within EU germplasm, thereby enabling the differentiation of genotypes. The established cut-off for determining identity status of the seed (i.e., self-pollinated or hybrid) was set at four or more heterozygous base calls.

### Plant phenotyping

The following traits were assessed in this study: *Stigma area and ovary diameter measurements* Image acquisition and annotation and quantification of carpel traits was performed as described in (17). The phases of stigma development were calculated based on the ± 15% of the maximum stigma area measured (also outlined in (17)).

#### Hybrid seed set assessment

To study seed set of the MS cultivars in a hybrid seed production setting, we manually counted the number of seeds and number of spikelets per spike. We assigned seeds to either apical, central, or basal sections based on their relative position along the spike. The apical section was defined as the top-most quarter of spikelets, the central section included spikelets from the central half of the spike, and the basal section consisted of the bottom-most quarter of spikelets. Additionally, we threshed spikes from the control plots and used seed counters (Seed Counter R-25 Plus and R-60 Plus, DATA Detection Technologies Ltd.) to determine seed number. In control plots, we calculated the grain number per spikelet by considering the average spikelet number per spike and the respective grain number per MS cultivar.

### Statistical analyses

To evaluate the effect of pollination time on grain per spikelet, we used a linear mixed model using a restricted maximum likelihood (REML) method from the R package lme4 v1.1.34 (20):

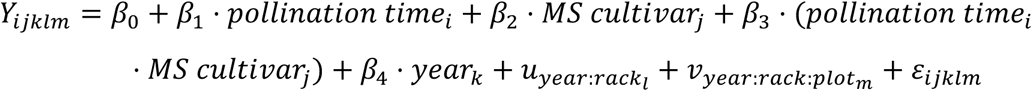

where *Y_ijklm_* is the observed grain number per spikelet value of the *i*th pollination time tested of the *j*th MS cultivar in the *k*th year in the *m*th plot of the *l*th rack. *u_year:rack_l__* is the random effect associated with the *l*th rack within the *k*th year. *V_year:rack:plot_m__* is the random effect associated with the *m*th plot within the *l*th rack and *k*th year. *ε_ijklm_* is the residual error term.

To account for variation between biological replicates we treated plot information as random effects. Diagnostic plots were generated to assess the goodness of fit of the model to the data (Supplementary Figure S5). R package lmerTest v3.1-3 was used to calculate *P* values (21).

The linear mixed model used to analyse the effect of pollination time on grain per spikelet per spike section is as follows:

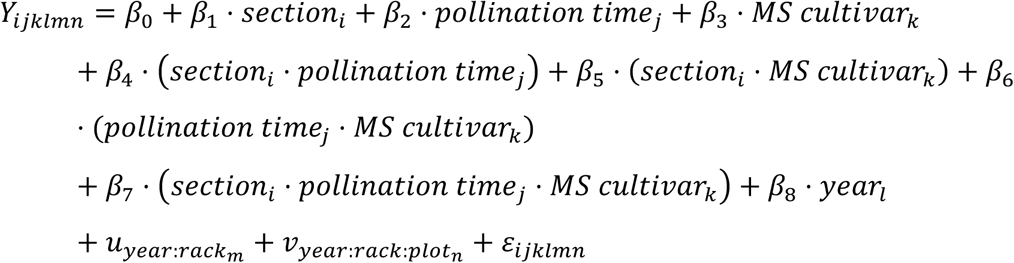

where *Y_ijklm_* is the observed grain number per spikelet value of the *i*th spike section on the *j*th pollination time tested of the *k*th MS cultivar in the *l*th year in the *n*th plot of the *m*th rack. Diagnostic plots are shown in Supplementary Figure S6.

To calculate the slopes for each combination of section and MS cultivar we used the “lm” function from the inbuilt R package stats v4.3.0 to fit the following linear model:

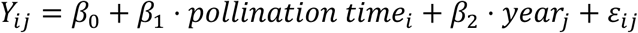

where *Y_ij_* is the observed grain number per spikelet value of the *i*th pollination time tested in the *j*th year for a particular section and MS cultivar.

## RESULTS

### Selection and characterisation of carpel development in the female parents

To understand how stigma longevity in the female parent influences hybrid seed set under field conditions, we first evaluated the patterns of stigma development in the absence of pollination in 29 MS cultivars derived from commercial inbred lines (provided by KWS and Syngenta) during the 2020 field season (Figure 1). We tracked the progression of stigmatic growth and deterioration at different timepoints, starting shortly after ear emergence at Waddington 9.5 (W9.5), and continuing at 3, 7, 13 and 18 days after W9.5 (DAW9.5). Hierarchical clustering (Figure 1A) and principal component analysis (PCA; Figure 1B) facilitated the identification of six distinct groups characterising diverse developmental patterns (see Supplementary Figure S7). Interestingly, PC1 seemed to explain the variation observed in the rate of stigma deterioration. For example, MS cultivars in groups 1, 2 and 3 showed slower progression of stigma deterioration (right section of PCA plot; Figure 1B and Supplementary Figure S7), while groups 5 and 6 represented the most prevalent developmental profile, including 20 of the tested MS cultivars (69%). Conversely, group 4, comprising cultivars 24500, 24522, and 24524, displayed a more premature stigmatic deterioration (left section of PCA plot; Figure 1B and Supplementary Figure S7).

**Figure 1.**
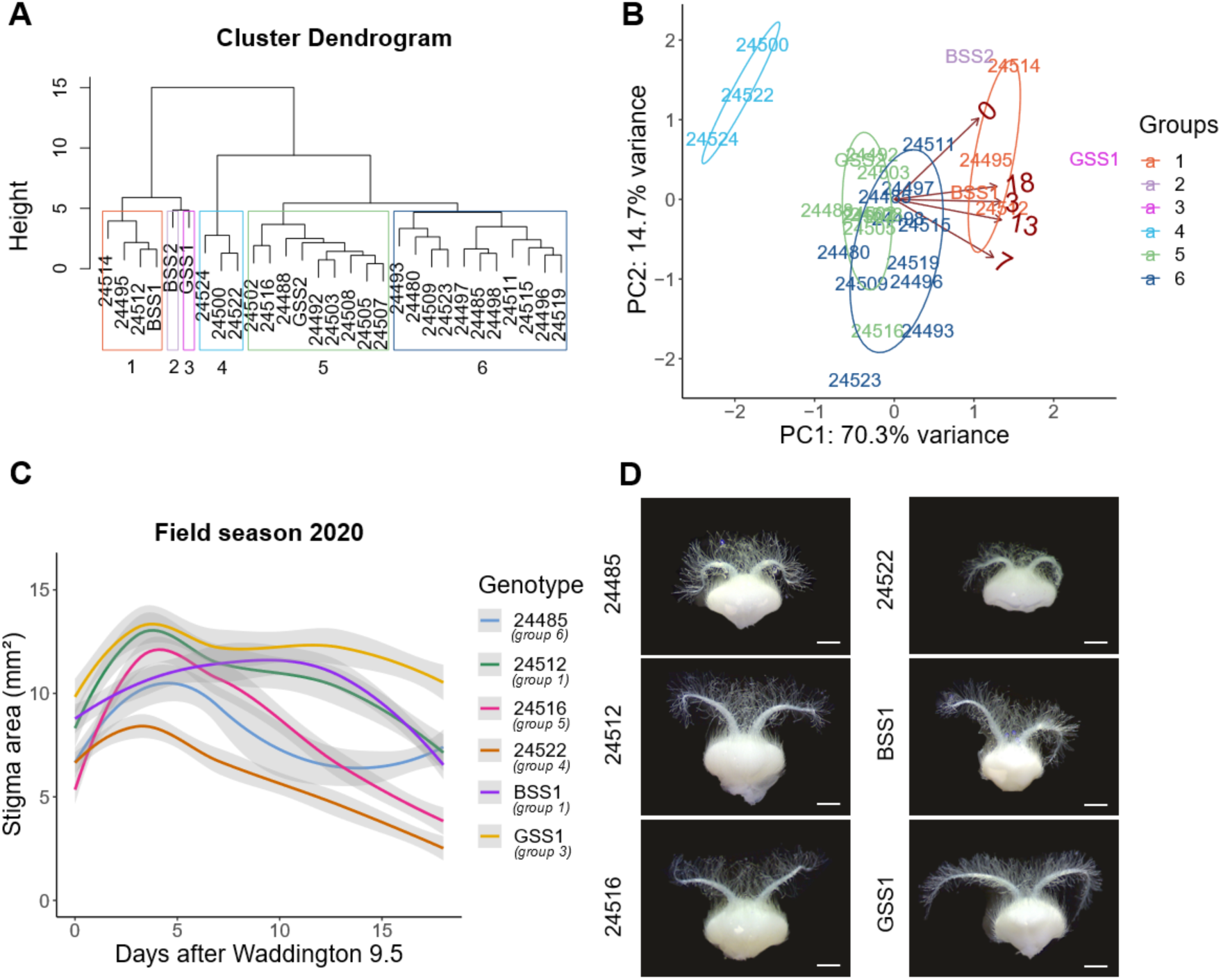
Characterisation of stigma traits for the selection of female parents. (A) Hierarchical clustering analysis to visualise the relationships between the 29 MS cultivars based on stigma area at 0, 3, 7, 13 and 18 DAW9.5. Coloured boxes define six distinct phenotypic groups labelled from 1 to 6. (B) Principal component analysis (PCA) of stigma area. MS cultivars are colour coded according to groups defined in (A), and dark red numbers refer to days after W9.5. (C) Developmental pattern of stigma area (mm^2^) for the selected MS cultivars (24485, 24512, 24516, 24522, BSS1 and GSS1) during the 2020 field season. Polynomial regression models at a 95% confidence interval (Loess smooth line) are shown. Grey shading represents the standard error of the mean (s.e.m). Five carpels from each of four plants were sampled at each timepoint. Note the colour scheme has been altered in relation to panels A and B. (D) Representative images of unpollinated carpels at 7 DAW9.5. Carpels are fixed in 95% ethanol and absolute acetic acid (75% v/v). Scale bar = 1 mm.

For each group, we chose a single MS cultivar to represent the observed variation, except for BSS2 in group 2, which was excluded due to its unusual growth and deterioration pattern (Supplementary Figure S7). Instead, from group 1, we selected two MS cultivars, namely 24512 and BSS1. These six cultivars collectively represent a substantial portion of the variability observed in stigma area throughout development (Figure 1C). Interestingly, these cultivars not only exhibited differences in the rate of deterioration, as exemplified by BSS1 and 24485, but also showed differences in stigma size, highlighted by a 5.5 mm² contrast at 7 DAW9.5 between GSS1 and 24522 (Figure 1D). These differences in stigma size might have important implications for the efficiency of hybrid seed production as they influence the level of stigma extrusion from the floret, thereby facilitating pollen capture (16,22,23). To ensure constant pollen flow throughout the hybrid seed production trials, we chose seven male fertile cultivars with good anther extrusion phenotypes and distinct flowering times. These cultivars were combined to form a mix of pollen donors for the 2021 and 2022 field seasons (see Materials and Methods). In 2021 and 2022, the pollination window spanned a period of 13 days. Notably, Nirvana was the earliest flowering cultivar, while the latest flowering cultivar varied between the two years (Supplementary Table S2). In some instances, flowering of the male line initiated prior to the MS cultivar reached W9.5, as observed with BSS1 and GSS1. In other cases, MS cultivars such as 24522 reached W9.5 before the earliest male cultivar, Nirvana, entered the anthesis phase. As a consequence, the effective pollination window differed among the MS cultivars.

To assess the carpel longevity of the selected MS cultivars during the 2021 and 2022 hybrid seed production trials, we replicated the phenotypic screenings carried out in 2020 to validate variations in stigma traits (Figure 2). During 2021 and 2022, alongside the evaluation of stigma traits, we quantified the diameter of the unfertilised ovary, given its potential role in floret opening and pollen access (14). To facilitate the analysis of stigma development, we used the method described previously for quantification of the growth, peak, and deterioration phases (Figure 2A and B) (17). Based on the onset of stigma deterioration, we identified three groups. Among the MS cultivars, 24522 was the earliest to enter the deterioration phase, occurring around 9 DAW9.5. Meanwhile, the stigmas of the MS cultivars 24516 and 24485 started to deteriorate around 12 DAW9.5. In contrast, GSS1, BSS1, and 24512 reached the deterioration phase roughly at 14 DAW9.5. Notably, BSS1 exhibited substantial variability between field seasons, with a 5-day difference in the onset of the deterioration phase (at 19 DAW9.5 in 2021 and at 13 DAW9.5 in 2022), consistent with previous data (17).

**Figure 2.**
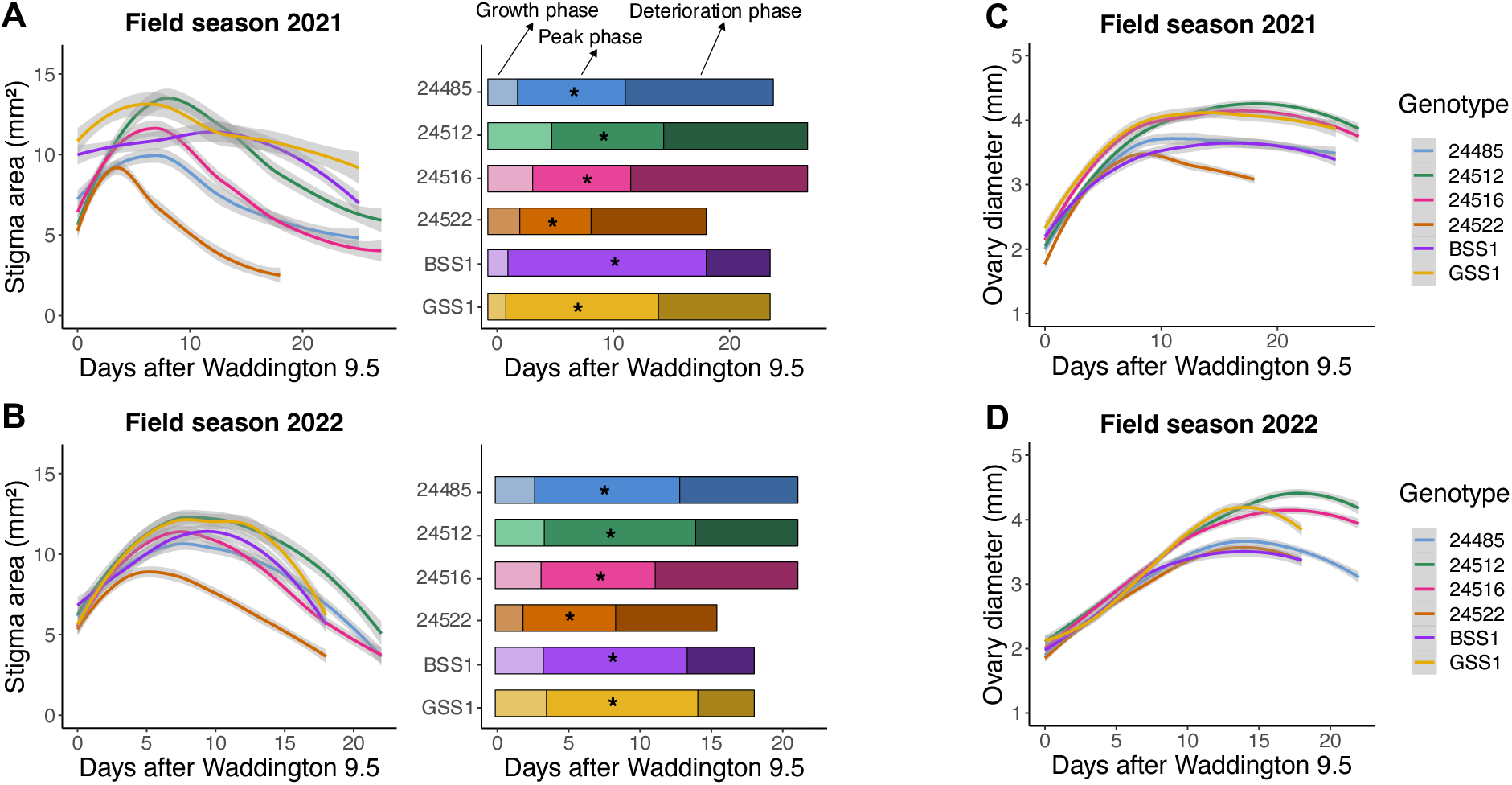
Phenotypic quantification of carpel development in the female parents across the 2021 and 2022 field seasons. (A, B) Developmental patterns of stigma area (mm^2^) and bar charts illustrating the growth, peak and deterioration phases represented in DAW9.5. The end of the deterioration phase is marked by the last sampling point and asterisks indicate days where stigma area peaked. (C, D) Developmental patterns of ovary diameter (mm). As for Figure 1C, polynomial regression models are shown for stigma area and ovary diameter. Five carpels from each of six plants were sampled at each timepoint.

Regarding stigma size, GSS1 and 24512 displayed the largest stigmas, reaching maximum areas of 13.5 mm^2^ in 2021 and 12.3 mm^2^ in 2022. An intermediate group, consisting of 24516, BSS1, and 24485, exhibited maximum sizes around 11 mm^2^. In contrast, 24522 had the smallest stigma, with a maximum size of 9 mm^2^.

Interestingly, we find that smaller stigmas reached their maximum size earlier than intermediate and larger stigmas, resulting in an overall 3-day difference in peak stigma area between the smallest and largest stigmas. Furthermore, we observed that 24522 consistently exhibited the smallest stigma size in both years, while growth patterns in the remaining cultivars exhibited some variation over the two years, with 2022 showing less variability between MS cultivars than in 2021 (Figure 2A and B).

In relation to ovary size, we observed two distinct groups: the first comprises GSS1, 24512, and 24516, which formed wider ovaries that reached a maximum diameter of 4.16 and 4.22 mm in 2021 and 2022, respectively. The second group exhibited narrower ovaries, with maximum widths of 3.56 and 3.57 mm in 2021 and 2022, respectively (Figure 2C and D). As seen for stigma size, this latter group reached its maximum ovary size, on average, 3 to 6 days earlier than the MS cultivars with broader ovaries, suggesting a potential correlation between carpel size and longevity. Additional analyses uncovered a positive correlation between carpel longevity and both stigma size (R = 0.6, *P* value = 0.0088) and ovary diameter (R = 0.23, *P* value = 0.36; Supplementary Figure S8). This result implies that larger carpels are associated with increased longevity, leading to delayed carpel deterioration. However, the latter correlation was weaker and not statistically significant, indicating that a larger sample size could better address the hypothesis regarding the potential association between carpel size and longevity.

### Evaluation of female performance in relation to hybrid seed set and stigma longevity

Following the assessment of stigma and ovary traits, we tested hybrid seed set performance of the six selected MS cultivars by allowing cross-pollination to happen during multiple time windows starting after W9.5 up to 10-15 DAW9.5 (Figure 3A). To control the timing of pollination, we used pollen proof barriers to prevent pollen flow between male donors and MS cultivars (Supplementary Figure S2 and S3). Cross-pollination was facilitated by removing the barriers, leading to progressively shorter pollination windows as the barriers were removed during the pollen-shedding period.

**Figure 3.**
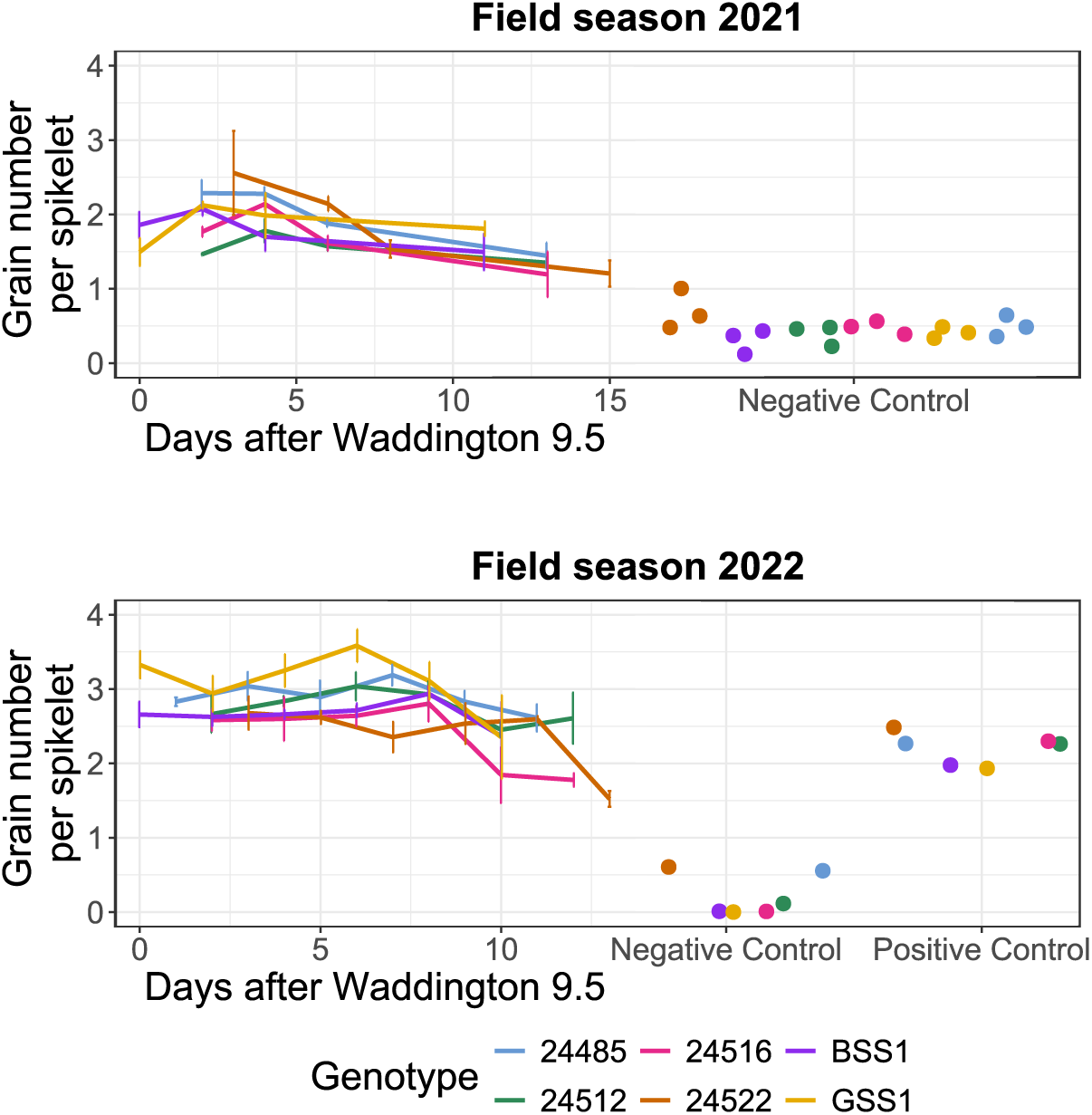
Temporal dynamics of hybrid seed yields. Line graphs illustrate the effect of pollination time on seed set (grain number per spikelet) for the different MS cultivars. Error bars denote the s.e.m. for each pollination window. In 2021 and 2022, four and six pollination windows were examined, respectively. Between five and ten spikes were sampled

Broadly, we observed substantial variation in hybrid seed set between 2021 and 2022, with a mean difference of one grain per spikelet greater in 2022 than in 2021 (0.94 ± 0.114; *P* value < 0.0001). Given that pollen dispersal is influenced by variations in humidity during the day, it is possible that the lower seed set in 2021 was caused by increased humidity brought on by precipitation and lower temperature for a significant portion of the time course (Supplementary Figure S4) (24). On average, hybrid seed set exhibited a slow yet statistically significant decline over time after W9.5, characterised by a negative rate of -0.054 ± 0.02 grains per spikelet DAW9.5^-1^ (*P* value = 0.007; Table 1). However, the presence of a significant “pollination date x MS cultivar” interaction term indicated that the timing of pollination had a non-uniform effect on seed set across the different MS cultivars. Specifically, MS cultivar 24522 was the only cultivar influenced by the timing of pollination, displaying a significant decline in hybrid seed set (linear slope = -0.056 ± 0.026 grains per spikelet DAW9.5^-1^; *P* value = 0.033; Table 1). Conversely, for the remaining MS cultivars, the rate at which seed set varied did not significantly differ from zero, indicating that seed set at later days was no different from earlier days (Table 1). Overall, our findings indicate that the duration of stigma receptivity or the longevity of the female parent has limited impact on the setting of hybrid seed in a free pollination experiment spanning ten to 15 DAW9.5.

**Table 1.**
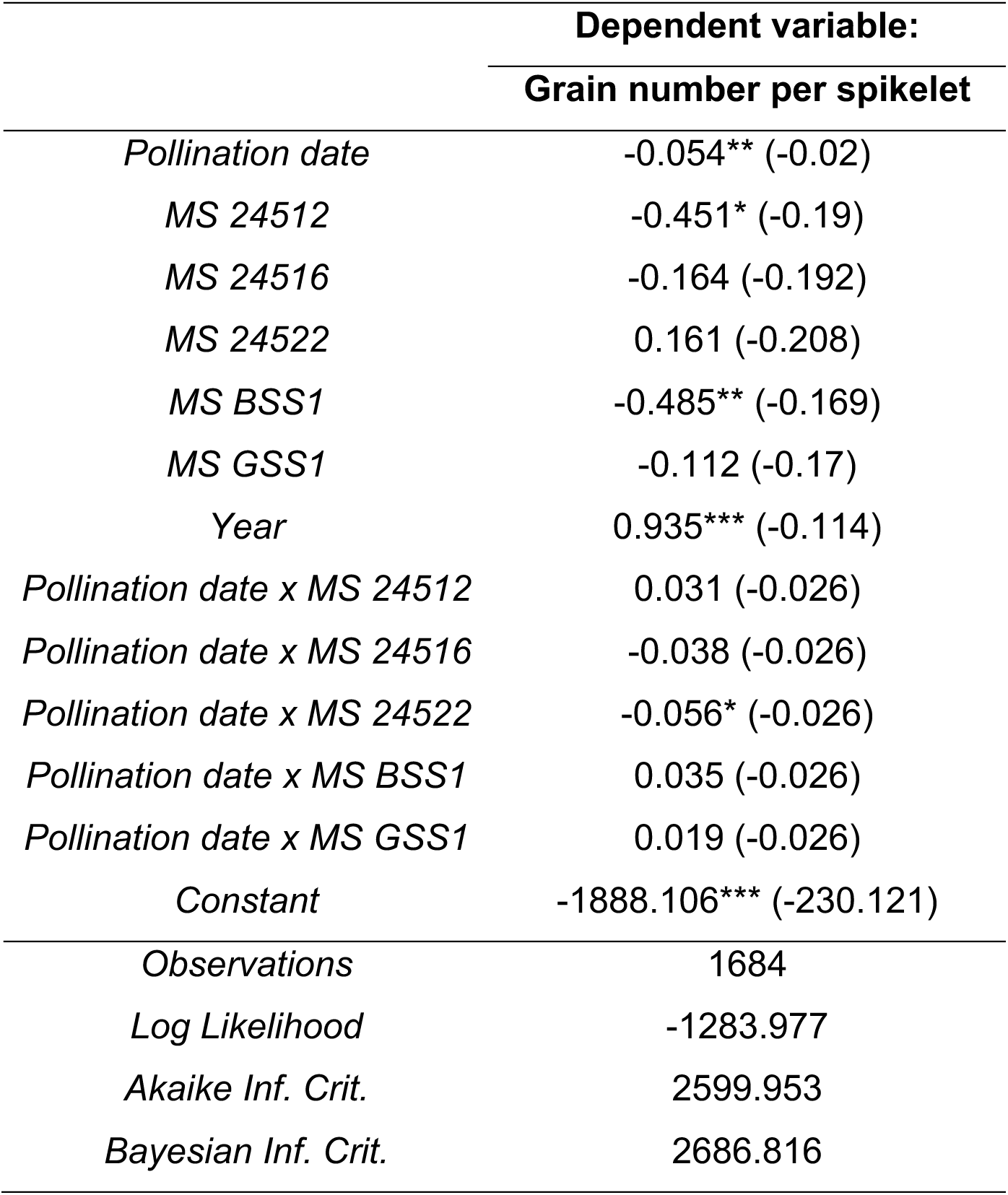
Regression table summarising linear mixed model results. Fixed effect coefficients along with their standard errors (in parenthesis) and P values are given for each relationship between the dependent variable (grain number per spikelet) and the independent variables. Key model fit statistics are also provided: Log Likelihood, Akaike Information Criterion (AIC), and Bayesian Information Criterion (BIC). A value of 0.05 indicates a decrease of one grain per spikelet in 20 days, such that a 20-spikelet spike will have 20 less grain after 20 days. *p<0.05; **p<0.01; ***p<0.001.

|  | **Dependent variable:** |
| --- | --- |
|  | **Grain number per spikelet** |
| *Pollination date* | -0.054** (-0.02) |
| *MS 24512* | -0.451* (-0.19) |
| *MS 24516* | -0.164 (-0.192) |
| *MS 24522* | 0.161 (-0.208) |
| *MS BSS1* | -0.485** (-0.169) |
| *MS GSS1* | -0.112 (-0.17) |
| *Year* | 0.935*** (-0.114) |
| *Pollination date x MS 24512* | 0.031 (-0.026) |
| *Pollination date x MS 24516* | -0.038 (-0.026) |
| *Pollination date x MS 24522* | -0.056* (-0.026) |
| *Pollination date x MS BSS1* | 0.035 (-0.026) |
| *Pollination date x MS GSS1* | 0.019 (-0.026) |
| *Constant* | -1888.106*** (-230.121) |
| *Observations* | 1684 |
| *Log Likelihood* | -1283.977 |
| *Akaike Inf. Crit.* | 2599.953 |
| *Bayesian Inf. Crit.* | 2686.816 |

Based on the maintained seed set rates across the complete sampling time course and the production of seeds in our negative controls where pollen barriers remained throughout the experiment, we asked if the seed formed were indeed hybrids. This question is justified by the challenges associated with achieving complete sterility in male sterility systems, particularly within the “BLue Aleurone (BLA)” system (25), which was used to develop MS cultivars 24485, 24512, 24516 and 24522. Notably, MS cultivars 24522 and 24485 appeared to be more significantly affected by residual fertility (Figure 3). To better understand how this might impact the interpretation of our results, we selected a random sample of the harvested F_1_ seeds from the test plots in 2021 and 2022 for hybridity assessment. While BLA-derived MS cultivars displayed the highest percentage of self-pollinated (i.e., homozygous) seeds, all hybridity levels significantly exceeded the 85% threshold required to grade seed set as high-quality hybrids (13). Thus, we conclude that hybrid seed production, and therefore, flower receptivity, remained effective across the various pollination windows tested, with the sole exception being a slight reduction in hybrid seed set observed in MS cultivar 24522, which is likely attributed to its premature stigma deterioration (Figure 2). This ability of MS cultivars to sustain seed set performance for more than 10 DAW9.5 is further exemplified by the positive controls implemented in 2022, where no pollen barriers were used and seed set levels closely mirrored those of plants pollinated at and after 10 DAW9.5 (Figure 3).

### Spatial distribution of hybrid seed set along the inflorescence

In this study, we focused exclusively on the developmental dynamics of carpels situated within central spikelets. However, it is well-known that maturation of the wheat florets (anthesis) begins in the middle of the spike and progresses bidirectionally towards the top and bottom of the spike, resulting in spike anthesis being spread over a two to six day period (26–28). This asynchronous pattern is mirrored in stigma development, with stigma from florets of central spikelets being more advanced than those at the apical and basal sections (Figure 4A and B).

**Figure 4.**
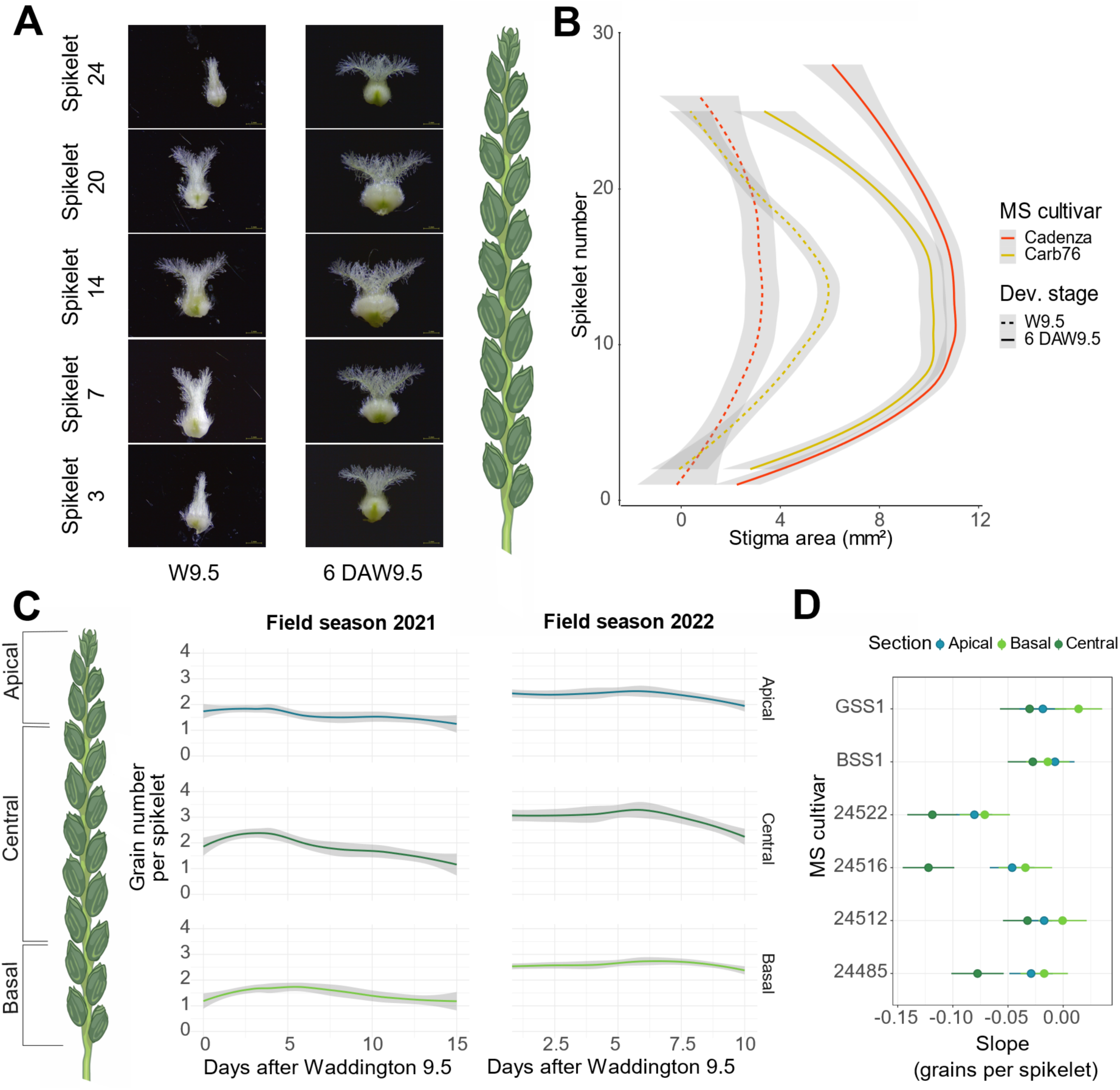
Bidirectional gradient of carpel development along the spike and distribution of hybrid seed set. (A) Representative images of carpels of MS cultivar Cadenza at Waddington 9.5 (W9.5) and at 6 days after W9.5 (DAW9.5). (B) Dynamics of stigma development along the spike for MS spring cultivars Cadenza and Carb76 under glasshouse conditions (n = 3). Polynomial regression models are shown for stigma area of florets 1 and 2. Red full line: Cadenza 6DAW9.5; red dotted line: Cadenza W9.5; yellow full line: Carb76 6DAW9.5; yellow dotted line: Carb76 W9.5. (C) Average association between hybrid seed set (grain number per spikelet) and the time of pollination per each section of the spike (i.e., apical, central, and basal) in 2021 and 2022. Polynomial regression models are shown. (D) Forest plot illustrates the progression of hybrid seed set (slope) for each section and MS cultivar. Horizontal lines represent the 95% confidence intervals.in each replication (n = 3). Negative and positive controls are depicted as single data points with n=3 for 2021 and n=1 for 2022.

To gain deeper insights into how these developmental gradients might have influenced both the temporal and spatial distribution of hybrid seed set, we dissected spikes into three sections: apical (top 25% spikelets), central (50%) and basal (bottom 25%; Figure 4C). By analysing the contribution of the different sections to hybrid seed production, we observed that grain number per spikelet was unevenly distributed along the spike, with central spikelets producing 0.5 and 0.4 grains per spikelet more than their apical and basal counterparts (*P* value < 0.0001). Spatial distribution of hybrid seeds therefore mimicked that of selfed seeds, where central spikelets produce the most grains (29,30). Moreover, we found that the temporal dynamics of grain number per spikelet differed between sections, and these dynamics were further influenced by the MS cultivar tested. To illustrate these complex interactions, we calculated the linear slopes for the temporal progression of grain number per spikelet within each section and MS cultivar combination while accounting for the variability observed across years (Figure 4D; see Materials and Methods for full models). Overall, we saw that hybrid seed production in apical and basal sections followed very similar trends, with slopes close to zero (Figure 4D).

This means that the number of grains formed in these apical and basal spikelets was consistently maintained over the sampled time, and hence, flowers were receptive to pollen across the complete time course. Notably, basal spikelets seemed to decrease at a slightly slower rate than apical spikelets, aligning with the progression of flowering (18), although the difference is minimal (linear slope apical section = - 0.03 ± 0.02; linear slope basal section = -0.02 ± 0.02). In contrast, central sections showed a more rapid decrease in hybrid seed set (linear slope: -0.07 ± 0.02), suggesting a significant decline at later pollination dates, likely explained by the gradual loss of flower receptivity as stigmas age. However, the lack of available pollen beyond these later dates did not allow for the extension of the time course to further investigate these trends. As anticipated by previous results (Table 1), not all MS cultivars behave similarly over time. For example, the reduction in hybrid seed set in MS cultivar 24522 can be attributed to a collective drop in all three sections as slopes are very similar to each other (linear slope apical section = -0.08 ± 0.02; central section = -0.12 ± 0.02; basal section = -0.07 ± 0.02). Additionally, we note that some MS cultivars showed a decline in seed set in the central section, while spikelets in apical and basal sections continued to perform equally well during the time course, as exemplified by 24516 and 24485. Finally, BSS1, GSS1 and 24512 displayed consistent seed set across all three sections, which aligns with a prolonged stigma lifespan (Figure 2A). In summary, we observe that hybrid seed production remains relatively stable in the apical and basal sections of the spike, whereas central spikelets exhibit a more rapid decline, likely due to their more advanced stigma developmental age. However, it is important to emphasise that these differences are minimal and the ability of the MS cultivars to sustain seed set performance over the sampled period is not significantly affected.

## DISCUSSION

Extending the duration of stigma receptivity in the female or male sterile (MS) parent has been commonly suggested as an important breeding target for improved hybrid seed production in crops (5,18,31). However, a noticeable knowledge gap exists in the literature, with very limited studies investigating the life cycle of the unpollinated stigma and its contribution to hybrid seed production. We attempted to address this gap by investigating how longevity of the wheat stigma in a MS parent influences out-crossing under field conditions when plants are exposed to free pollination.

Our analysis of carpel development of MS cultivars showed that, in the absence of pollination, the stigma survives between 9 and 14 days after stigmatic branches begin to unfold. From the subset of MS cultivars selected, we found that the majority entered the phase of stigma deterioration between 12 and 14 DAW9.5; with the sole exception of 24522, which initiated stigma deterioration as early as 9 DAW9.5 (Figure 1A and B). These MS cultivars also showed differences in the magnitude of carpel growth. Interestingly, larger stigmas and wider ovaries were mostly linked to MS cultivars featuring delayed carpel deterioration, such as 24512 and GSS1. On the contrary, those with more premature deterioration, like 24522 and 24485, displayed smaller carpels. These initial observations prompted questions about a potential association between carpel size and longevity, which was partially substantiated by additional correlation analyses (Supplementary Figure S8). While it is premature to speculate on the biological implications of these findings, it is reasonable to hypothesise that the selection of larger carpels – already a target for improved pollen access due to larger stigmatic area and wider ovary (12,14) – may also lead to indirect selection of carpels that survive for longer periods. Nonetheless, further research into understanding this link and its effect on hybrid seed set is needed.

The scarcity of studies investigating the role of stigma receptivity on cross-pollination and hybrid seed production can be attributed to the myriad of challenges inherent in conducting such experiments in the field. Wind-pollination relies on the range and quantity of pollen dispersal, which is greatly influenced by factors like wind direction and speed (downwind vs. crosswind), sunlight, temperature, precipitation levels, and relative humidity during anthesis (9,18). This context helps explain the significant increase in average hybrid seed yield observed in 2022 compared to the preceding year, when lower temperatures and higher precipitation were reported (Figure 3 and Supplementary Figure S4). Such fluctuations in hybrid seed yields across field seasons are not uncommon and have been documented in other studies, where environmental factors contribute up to 65% of the total variation in hybrid seed yield (13,32). Another challenge stems from the limited availability of late flowering pollinators. This constraint made it impractical to study hybrid seed set performance of female parents during the deterioration phase as fresh pollen became unavailable beyond the 10 to 15 DAW9.5 pollination window (Figure 3). Alternatively, different sowing dates could be used to alleviate this limitation, allowing for a more extended study period. Additionally, other factors that have not been considered in this study, and that could significantly influence cross-pollination and hybrid seed set, such as variability in pollen accessibility due to floret openness, deserve further exploration (5).

Despite all the challenges presented by open pollination experiments, we found a remarkable consistency in the patterns of hybrid seed set across MS cultivars and field seasons (Figure 3). Surprisingly, there was no substantial reduction in hybrid seed set during the 10 to 15 day pollination time course, suggesting sustained stigma receptivity up to 10 days after stigmatic branches began to unfold (i.e., shortly after ear emergence). Due to the lack of viable pollen at later time points, it was not possible to extend the time course to study the reproductive potential of the stigma during the deterioration phase; therefore, this hypothesis remains untested.

Nonetheless, we showed that the overall absence of reduced hybrid seed set is partially masked by asynchronous nature of stigma development along the spike (Figure 4), with central spikelets displaying a slightly greater drop in seed set compared to younger spikelets in apical and basal regions of the spike. Despite the biological relevance of understanding the dynamics of hybrid seed set during the latest stages of stigma development, from an agronomic standpoint, it is of minor significance. During the developmental stage of commercial hybrid programs (i.e., test crosses), pollen release typically occurs within a 10-day window, ranging from five days before ear emergence of the female parent (approximately W9.5) to five days after ear emergence (Supplementary Figure S9) (32). If we take this timeframe into account, we can conclude that stigma receptivity does not represent a major barrier in hybrid seed production, as female parents are able to sustain similar levels of seed set up to 10 DAW9.5. These findings suggest that future breeding efforts aimed at enhancing cross-pollination efficiencies could benefit from considering other desirable qualities of the female parent, such as ovary and stigma sizes, as well as floret openness. In rice, for instance, the percentage of exserted stigmas, largely determined by stigma size, correlates positively with hybrid seed set, and numerous quantitative trait loci have been identified that associate with this trait (33,34). Recent research in bread and durum wheat has revealed significant genotypic variation in stigma length; however, its contribution to hybrid seed set remains unexplored (16). Thus, a comprehensive analysis of these traits is essential to collectively determine the most critical factors for future breeding efforts which will then lay the groundwork for subsequent molecular studies. Understanding the biology of the wheat stigma will undoubtedly provide new opportunities for improving the cost-effectiveness of hybrid seed production.

## Acknowledgements

The authors would like to thank Marco Fioratti for assisting with the statistical analysis for the progression of hybrid seed set.

## Author contributions

M.M-B, N.B., Y.M., C.U., S.A.B. conceptualization; M.M-B., J.S. investigation; M.M-B., C.U., S.A.B. formal analysis; N.B., Y.M. resources; M.M-B. writing – original draft; M.M-B, N.B., Y.M., C.U., S.A.B. writing – review and editing; C.U., S.A.B. supervision.

