## Supplemental Figures for "Stigma longevity is not a major limiting factor in hybrid wheat seed production"

#### Supplementary Tables

**Supplementary Table S1. Plant scores of pollen donors selected for 2021 and 2022 hybrid seed production field trials.** Data obtained from KWS based on a 2020 field trial. AEX: anther extrusion (0 = no anthers extruded; 3 = maximum anther extrusion); EE: ear emergence.

| Pollen donor | AEX | EE | Height (cm) |
|---|---|---|---|
| Nirvana | 2.26 | 24/05/2020 | 70 |
| Elysee | 2.93 | 27/05/2020 | 100 |
| Quartz | 2.4 | 29/05/2020 | 56 |
| Stava | 1.68 | 03/06/2020 | 95 |
| Creator | 2.78 | 03/06/2020 | 64 |
| Poros | 2.33 | 04/06/2020 | 109 |
| Piko | - | - | - |

Note: No data was available for Piko based on the KWS 2020 field trial. Piko was selected for its excellent male floral traits (e.g., anther extrusion, height; Boeven et al., 2018). It also shows late ear emergence compared to most of pollen donors used in UK pollination trials.

**Supplementary Table S2. Flowering time (anthesis) of pollen donors in 2021 and 2022.** Flowering time was assessed in test plots without mix cultivation. Note that not all the pollen donors were grown separately in each of the two seasons.

| Pollen donor | Anthesis (2021) | Anthesis (2022) |
|---|---|---|
| Nirvana | 11/06/2021 | 26/05/2022 |
| Elysee | 15/06/2021 | 05/06/2022 |
| Quartz | - | 02/06/2022 |
| Stava | - | 07/06/2022 |
| Creator | - | 08/06/2022 |
| Poros | 22/06/2021 | - |
| Piko | 24/06/2021 | 03/06/2022 |

### Supplementary Figures

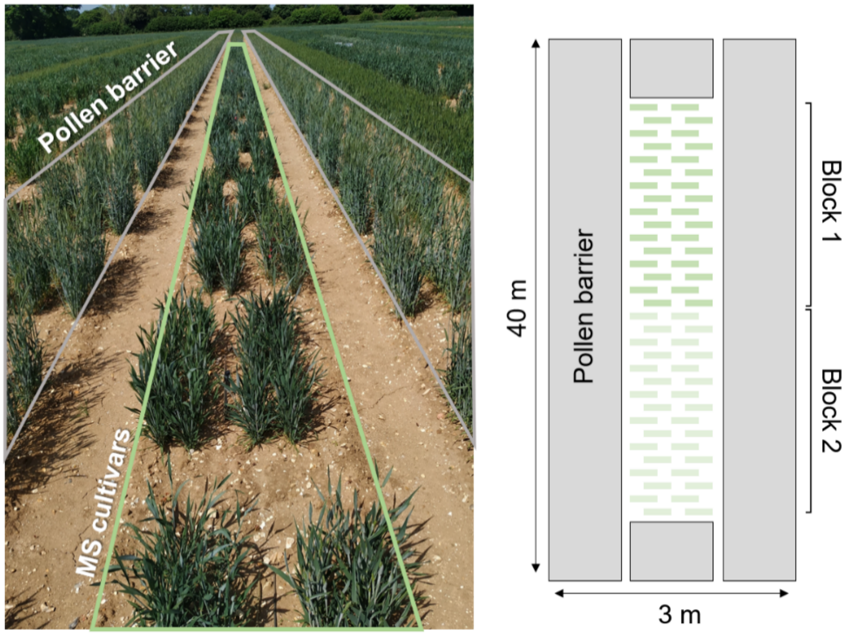


**Supplementary Figure S1. Visual and schematic representation of the field layout in 2020.** MS cultivars (in green) were grown surrounded by a continuous sowing of sterile rye that was used as pollen barrier. Plots were replicated twice in 2020 (n = 29 per block). This layout aligns with experiments focused on characterising the development of unpollinated carpels and was replicated in both 2021 and 2022.

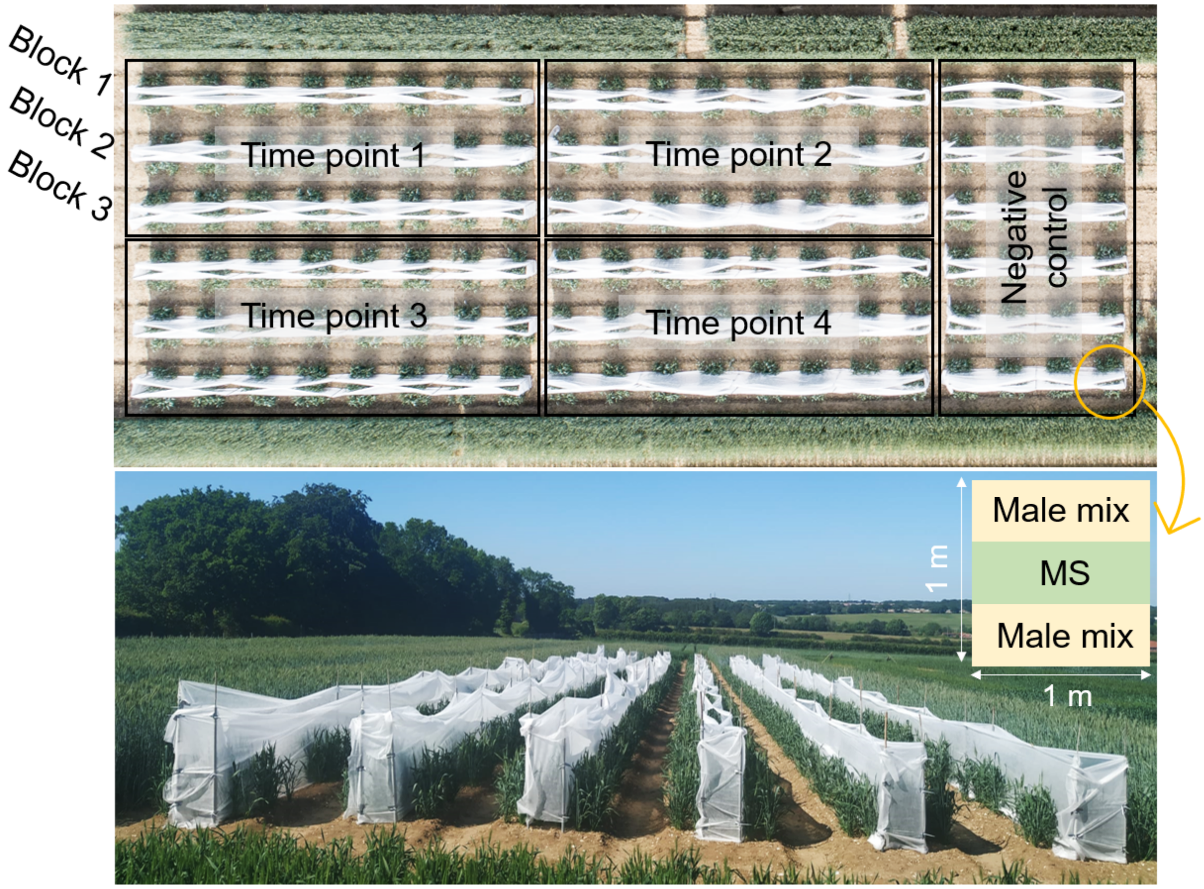


**Supplementary Figure S2. Field trial layout of the 2021 hybrid seed production trial before flowering.** Isolation walls encircle the MS rows, blocking pollen flow from adjacent male fertile plants. Note that the setup required the complete removal of these walls for each block.

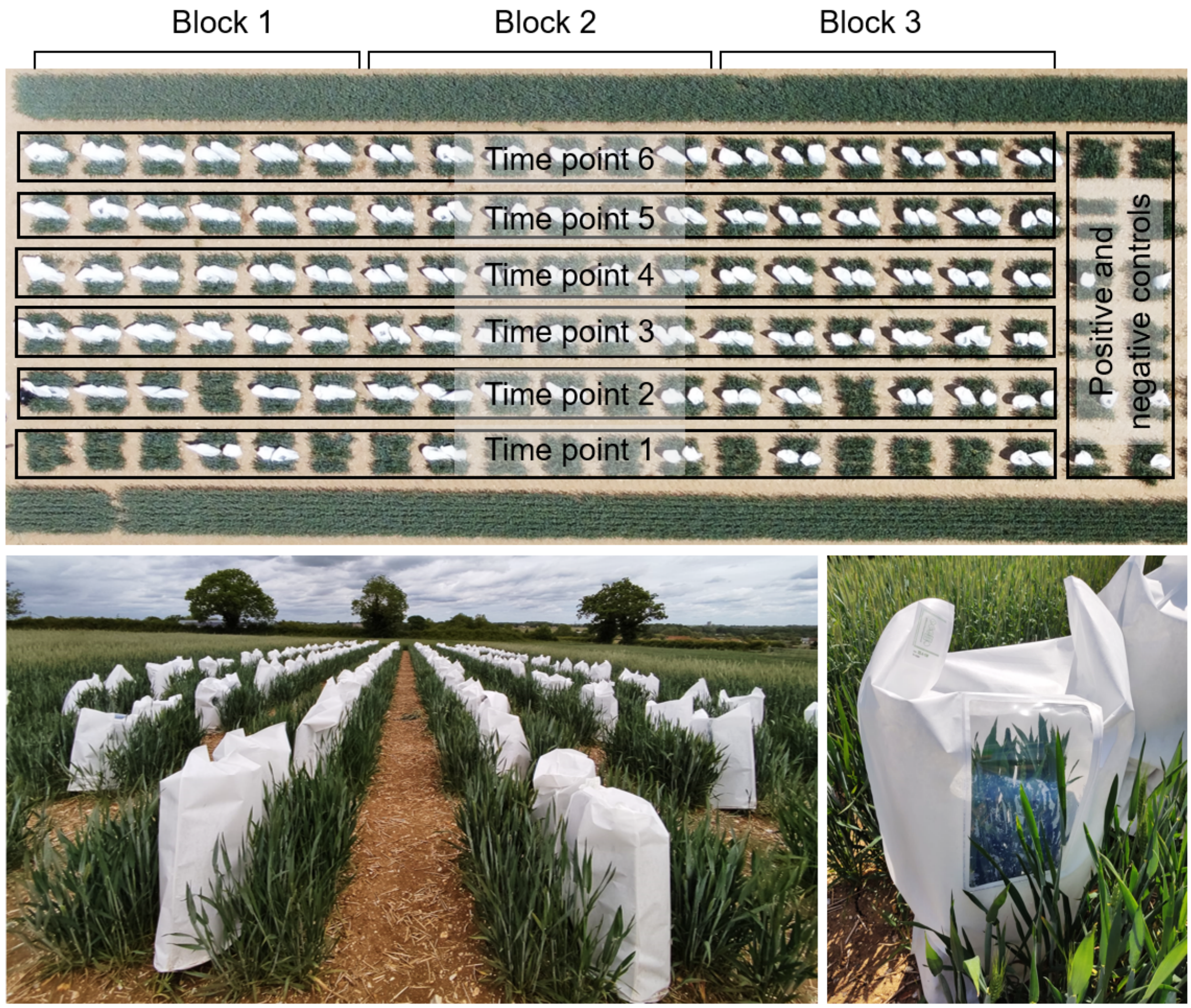


**Supplementary Figure S3. Field trial layout of the 2022 hybrid seed production trial before flowering.** Pollination bags covering the MS rows, blocking pollen flow from adjacent male fertile plants. Note that some bags were already removed at time point 1 and 2 as this setup, contrary to 2021, allowed for the individual removal of bags.

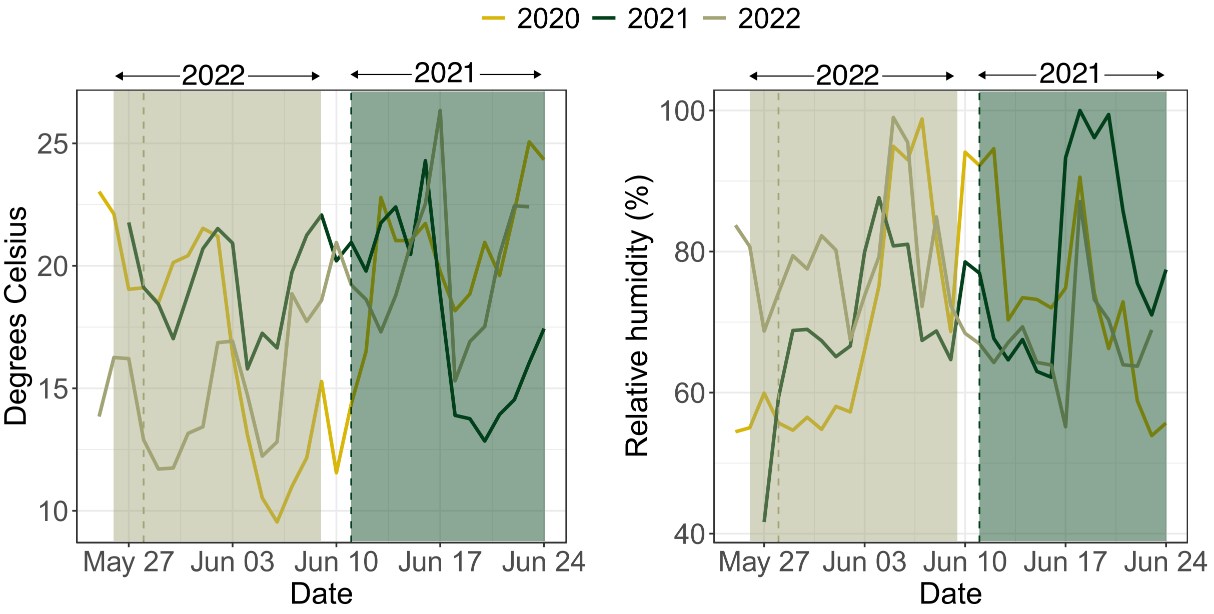


**Supplementary Figure S4. Environmental conditions recorded during 2020, 2021 and 2022 field seasons.** Left panel illustrates mean daily temperatures in degrees Celsius recorded during the field experiments. Shaded rectangles indicate the beginning and end of 2021 and 2022 time courses (i.e., pollination windows). Average W9.5 dates are indicated by dashed lines for 2021 and 2022 hybrid seed set time courses. Right panel shows the water vapor contained in the air expressed in percentage of relative humidity.

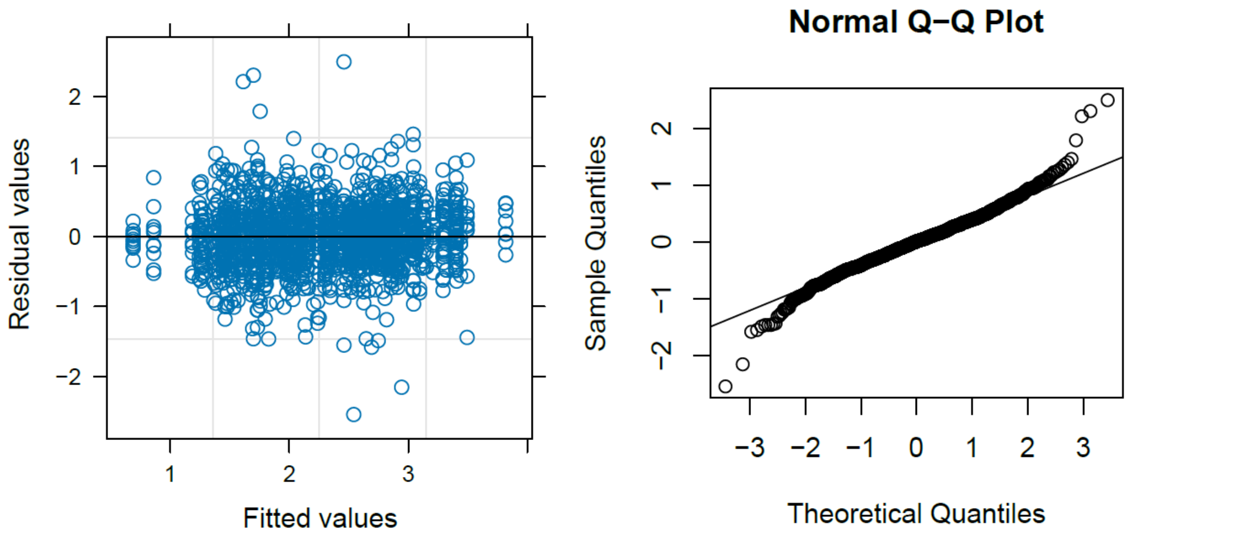


**Supplementary Figure S5. Diagnostic plots of residual values.** Scatter plot on the left shows the variance of the Pearson's residuals is constant with a fairly uniform distribution of the residuals indicative of a well fitted model. On the right, the quantile plot does not raise significant concerns around normality of the residuals.

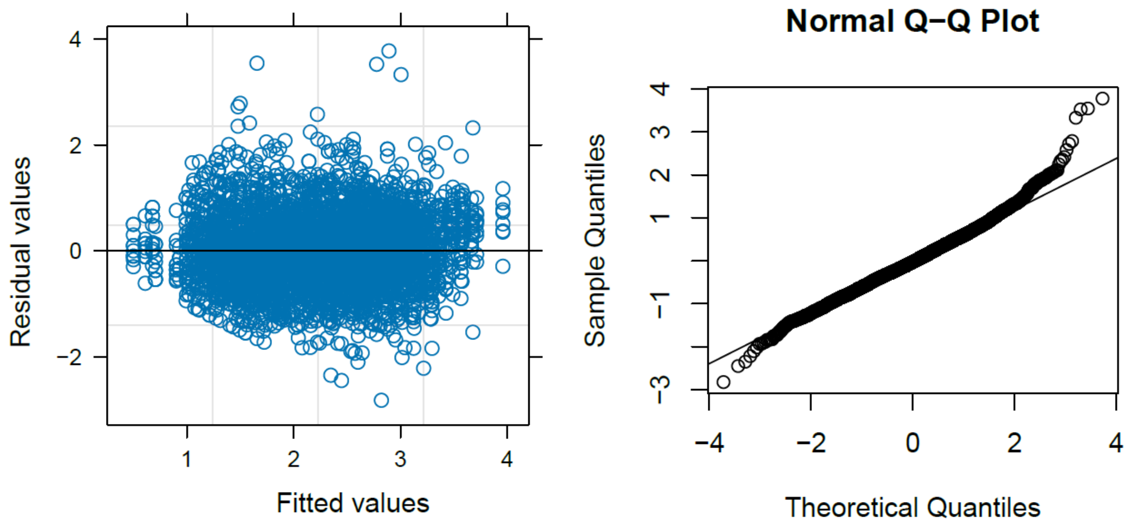


**Supplementary Figure S6. Diagnostic plots of residual values.** Scatter plot on the left shows the variance of the Pearson's residuals is constant with a fairly uniform distribution of the residuals indicative of a well fitted model. On the right, the quantile plot does not raise significant concerns around normality of the residuals.

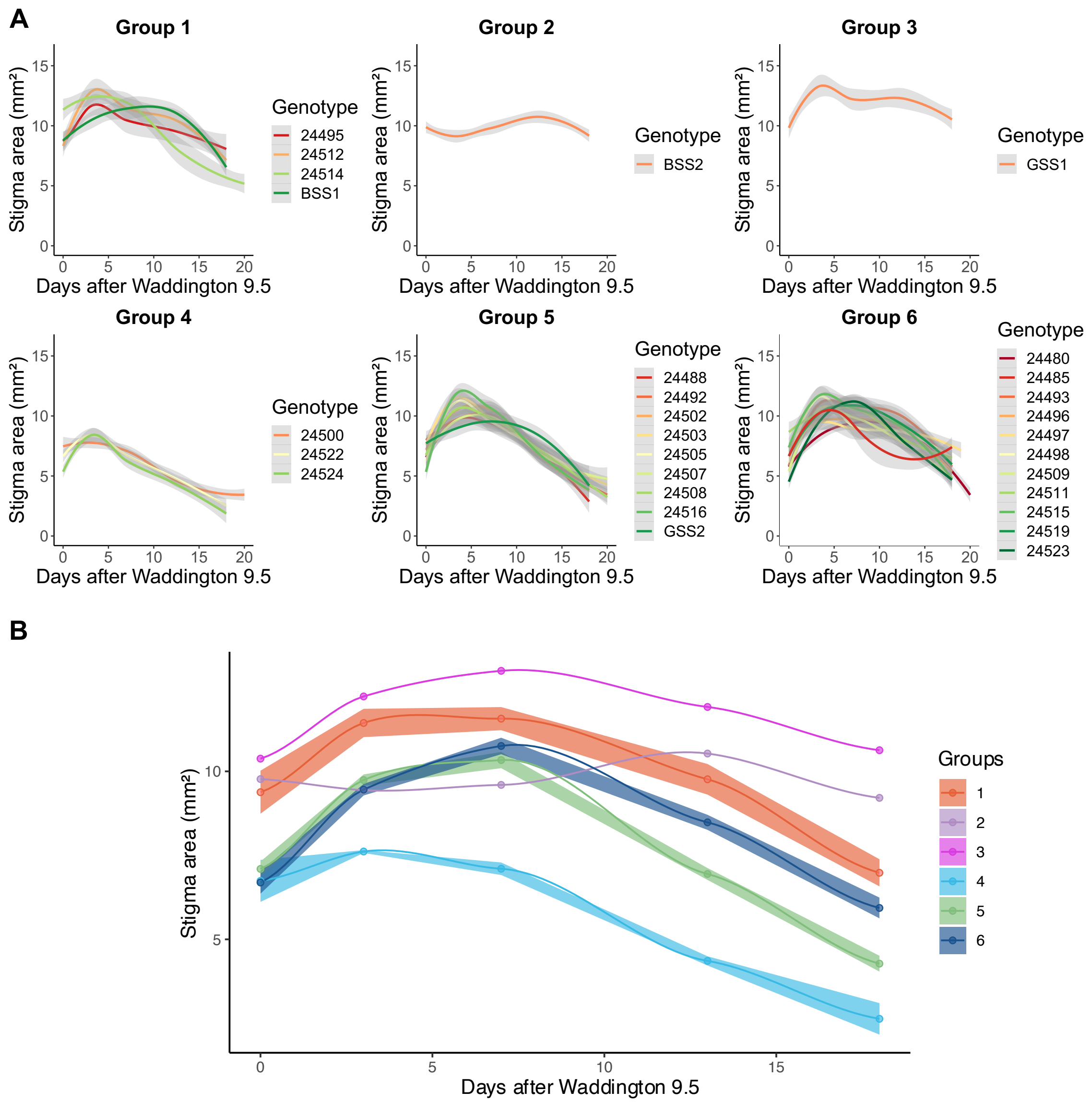


**Supplementary Figure S7. Phenotypic diversity observed for stigma development during the 2020 field season.** (A) Each plot represents a distinct group based on clustering analysis (see **Error! Reference source not found.**A). Polynomial regression models are shown for stigma area. Grey shading represents the standard error of the mean (s.e.m). Five carpels from each of four plants were sampled at each timepoint. (B) Ribbon plot illustrates how stigma area ($mm^2$) varies over the range of the six distinct groups. The width of each ribbon represents s.e.m.

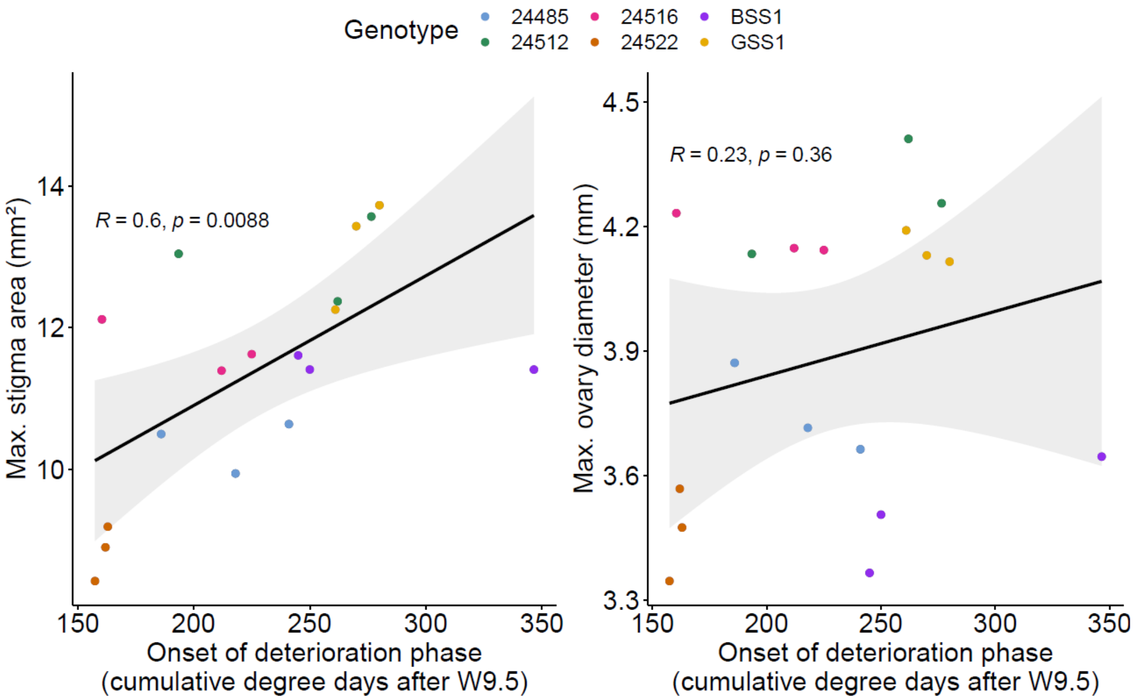


**Supplementary Figure S8. Relationship between carpel longevity and carpel size.** Scatter plots display the relationship between the onset of deterioration phase and maximum stigma size (left) and maximum ovary diameter (right) for all six MS cultivars selected. The onset of the deterioration phase is measured in cumulative degree days after W9.5 to accommodate variation in temperatures between field seasons 2020, 2021 and 2022 (n = 3). Regression lines are shown along with Pearson correlation coefficients and significance levels. Grey shaded area represents confidence intervals for the regression lines.

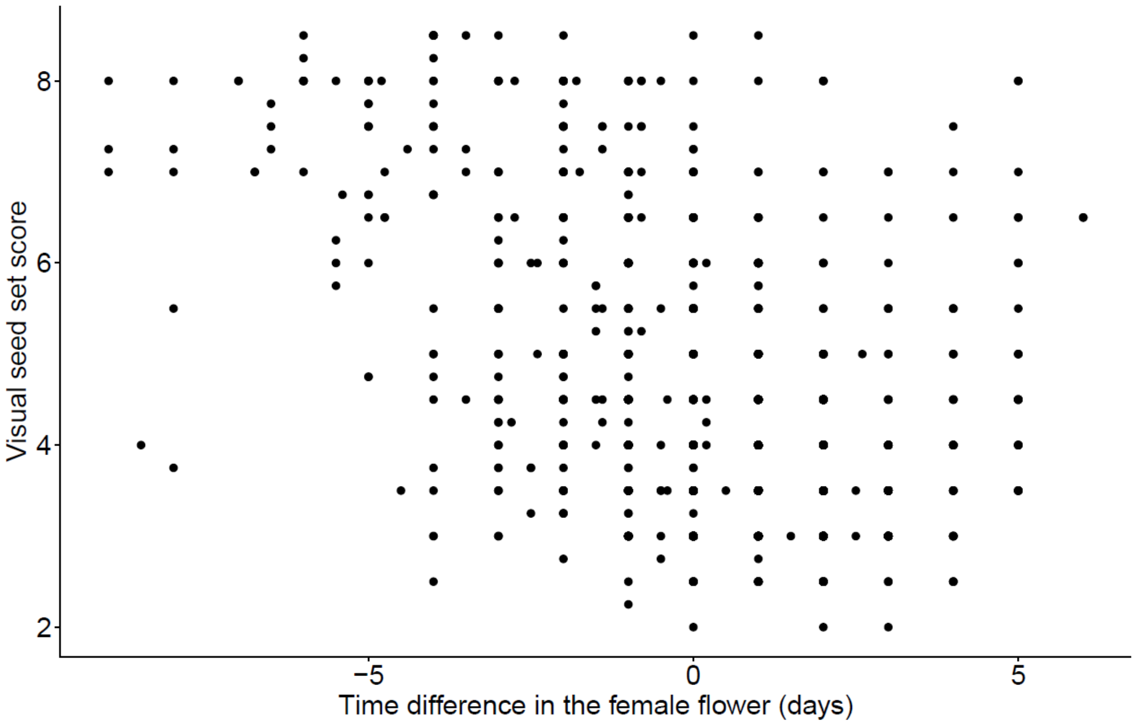


**Supplementary Figure S9. Visual seed set scores for the 2,428 unique combinations.**

Visual seed set scores range from 1 (seed set similar to a fully fertile spike) to 9 (no seed set). X-axis indicates difference in flowering dates between the female parent and pollen donor. Negative values represent hybrid combinations where anthesis of the pollen donor started “n” days before full ear emergence of the female; 0 indicates that pollination coincides with ear emergence of the female parent; and positive values indicate pollination started “n” days after full ear emergence of the female.
